# Robust and tunable performance of a cell-free biosensor encapsulated in lipid vesicles

**DOI:** 10.1101/2022.03.02.482665

**Authors:** Margrethe A. Boyd, Walter Thavarajah, Julius B. Lucks, Neha P. Kamat

## Abstract

Cell-free systems have enabled the development of genetically encoded biosensors to detect a range of environmental and biological targets. By encapsulating these systems in synthetic membranes, we can reintroduce features of the cell membrane, including molecular containment and selective permeability, which together could significantly enhance sensing capabilities. Here, we demonstrate robust and tunable performance of an encapsulated fluoride riboswitch inside of lipid vesicles. The riboswitch remains active upon encapsulation in lipid vesicles and responds to fluoride added to the surrounding solution. We find the sensitivity of the encapsulated sensor can be tuned by varying membrane composition. We then show that encapsulation protects the sensor from degradation by the sample and use two types of genetically encoded outputs to detect fluoride in real-world samples. This work establishes the feasibility of vesicle-encapsulated cell-free systems to detect environmentally relevant small molecules.

## Introduction

Cell-free systems have emerged as a powerful technology to detect a wide variety of molecular signals, including chemical contaminants relevant to the environment and human health (1–9) and markers of disease and infection (10–17). By reconstituting purified cellular machinery *in vitro*, these systems enable use of natural microbial sensing mechanisms in a low-cost, distributable, and easily tunable platform. Despite these key advantages, removal from the cell also eliminates certain features of the cell’s native membrane barrier - such as reaction containment, protection from reaction inhibitors, and selective gating - all of which can add important functionality to cell-free biosensors (18).

Efforts to deploy sensors highlight these limitations caused by the absence of cellular membranes. For example, without a barrier between the sensor and the sample, detecting targets in complex matrices like polluted water or biological samples requires additional modifications to the reaction or preparation protocols (6, 19, 20). Cell-free sensors are also sensitive to dilution, and therefore require a controlled reaction environment (21). One strategy to mitigate these limitations is to recapitulate some of the lost features of the cell membrane by encapsulating cell-free sensors inside of synthetic membranes.

Encapsulation enables tuning of the reaction environment on a molecular scale, enabling control of molecular interactions and addition of active membrane features to advance sensing capabilities, all while maintaining many of the tunable, advantageous features of cell-free systems (18).

There are two major considerations in designing encapsulated cell-free sensors: determining the impacts of a confined reaction environment on sensor function and choosing an appropriate target molecule and application. In terms of reaction confinement, the small scale of the encapsulated environment can impact reactant loading, reaction time, and limit of detection (22–24). These effects have been shown to impact the basic processes of gene expression (25), which in turn affects cell-free biosensors that regulate reporter gene expression at the level of transcription or translation (26). Of the wide range of genetic regulatory networks used for biosensing, RNA-based biosensors that regulate transcription require the fewest components and operate on a faster timescale (3, 27), which may reduce the impacts of confinement on sensor function. Riboswitches — noncoding RNA elements upstream of protein coding genes that change their structure in response to specific ligands to regulate gene expression — could offer an opportunity to address these constraints due to reaction confinement.

Previous proof-of-concept studies have focused on encapsulation of two synthetic, translationally regulated riboswitches that respond to membrane-permeable signals: theophylline (21, 28, 29) and histamine (30). Both riboswitches have been successfully encapsulated in bilayer vesicles, generating either a fluorescent protein readout or a protein-mediated response upon analyte entry into the vesicle interior (21, 28–30). Encapsulation of transcriptionally regulated riboswitches has proven difficult to date, however; efforts to encapsulate a transcriptionally regulated adenine riboswitch showed poor switching activity and were subsequently abandoned (29). This could be due to specific features of the adenine riboswitch or due to a general property of transcriptional riboswitches, which require dynamic conformational changes during transcription to enact their mechanism – a process which could be impacted by general features of confinement or electrostatic interactions with the lipid bilayers (31, 32). Despite these potential challenges, the mechanisms underlying transcriptionally regulated riboswitches are being further uncovered (33). These sensors have demonstrated the feasibility of detecting environmentally important analytes in cell-free systems and can function with RNA-level outputs (3) - a key feature which may mitigate resource constraints - motivating further efforts for their encapsulation and deployment.

A second major consideration in encapsulated sensor development is the selection of an appropriate target and application. Of the many potential uses of encapsulated biosensors, water quality monitoring is one of the most compelling from a global perspective. One in three people globally lack access to safe drinking water (34), and the ability to identify contaminated water sources is essential for their quarantine or remediation (35). Fluoride is among the most concerning of these contaminants; chronic exposure to fluoride binds it to the calcium in teeth and bones, weakening them and causing lifelong health consequences (36). From both environmental and anthropogenic sources, fluoride exposure is especially problematic in parts of China, Africa, South America, and India (36, 37), with high fluoride concentrations also found in groundwater across the United States (37). This diversity of sample sources comes with a corresponding increase in potential reaction inhibitors, presenting the need for a robust sensor that retains function in complex matrices. Encapsulated fluoride biosensing reactions would address this need, delivering far-reaching global health benefits and establishing a framework to address future water quality challenges.

In this study, we sought to develop vesicle-based sensors for fluoride by encapsulating a transcriptionally regulated, fluoride-responsive riboswitch within bilayer membranes (Figure 1). We first encapsulate the riboswitch, then demonstrate its ability to detect externally added fluoride and show that membrane composition can be modified to tune sensitivity to exogenous ions. We also demonstrate that encapsulation protects cell-free reactions from sample degradation, particularly from extravesicular degradative enzymes. Finally, we couple riboswitch output to both fluorescent and colorimetric reporters and show that vesicle-based sensors can detect fluoride in real-world water samples. This work demonstrates the potential of encapsulated, riboswitch-based sensors for biosensing applications, complimenting existing cell-free sensor engineering strategies and enabling sensing in otherwise inhospitable environments.

**Fig. 1.**
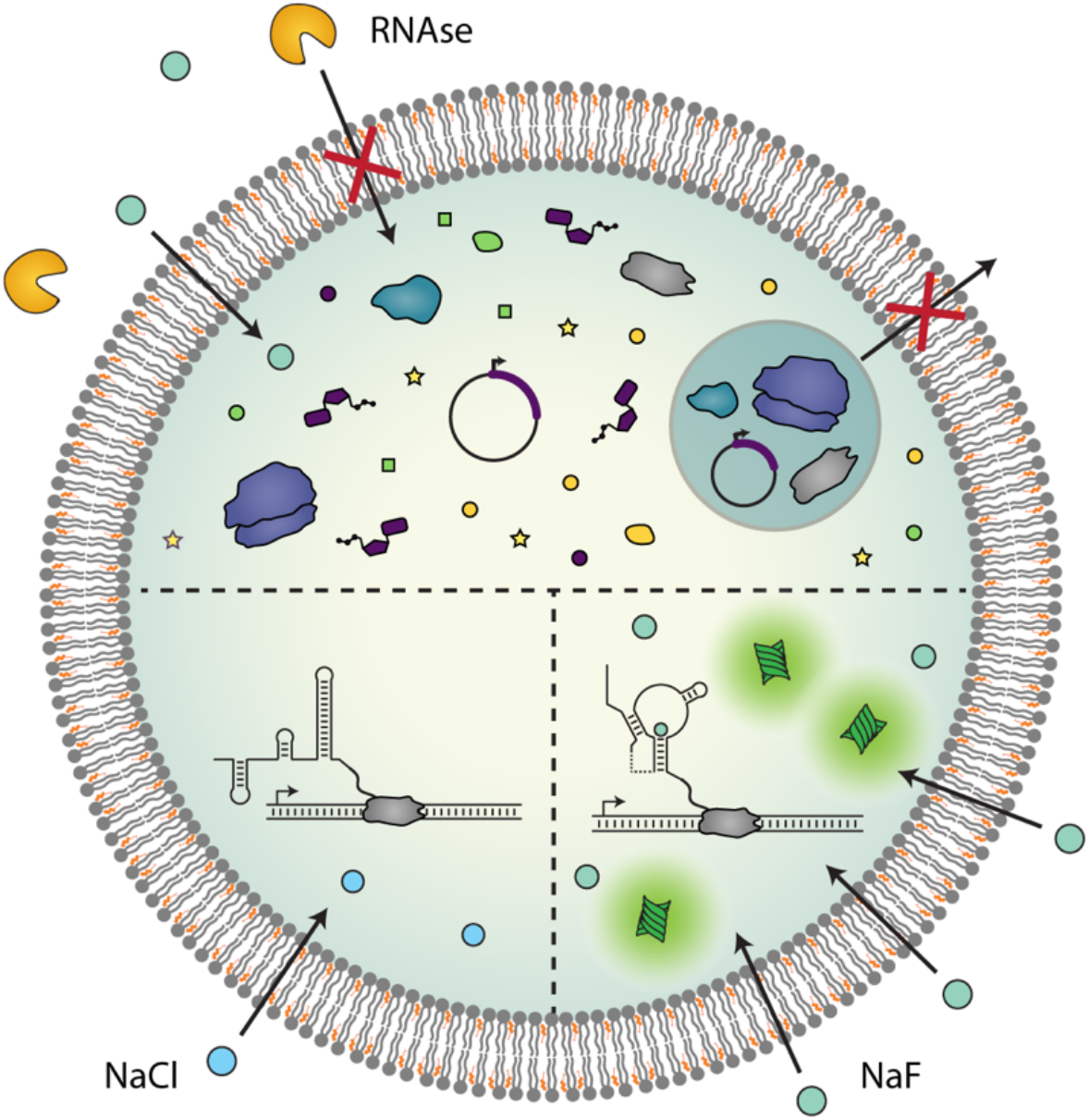
Encapsulated cell-free sensors. Encapsulation of cell-free systems creates a semipermeable barrier between sensor components and the environment, which modulates their molecular interactions. Reactants are contained within the vesicle interior, while proteins and other large molecules in the external sample are excluded from vesicle entry (top). Small, membrane-permeable molecules can diffuse into the vesicle interior, initiating a riboswitch-mediated response that is specific to an analyte of interest (bottom right). The riboswitch folds into a terminating conformation in the absence of sufficient concentrations of target analyte (bottom left).

## Results

### A transcriptionally regulated fluoride riboswitch can function inside lipid vesicles

We first sought to confirm that a transcriptional riboswitch can function when encapsulated inside lipid vesicles. For the riboswitch, we chose the fluoride responsive riboswitch from *Bacillus cereus*, which we previously showed can be used to control the expression of several different reporter proteins and fluorescent RNA aptamers in bulk *E. coli* extract-based cell-free systems (3). In this system, the fluoride riboswitch is encoded within a single DNA template, downstream of a consensus *E. coli* promoter sequence, and upstream of a reporter coding sequence. In the absence of fluoride, *E. coli* polymerase transcribes the riboswitch sequence, causing it to fold into a conformation that exposes a transcriptional terminator hairpin and subsequently causes RNA polymerase to stop transcription (38). In the presence of fluoride, fluoride binding to the riboswitch aptamer domain prevents the terminator from folding, allowing transcriptional elongation of the reporter coding sequence.

In this study, we first chose to use a super folder green fluorescent protein (GFP) reporter, as it allows convenient measurement of riboswitch activity. For the cell-free system, we used an *E. coli* S30 lysate prepared with runoff and dialysis, which has been shown to allow the function of biosensors that require bacterial polymerases (39). Embedding the riboswitch DNA template into the extract system alongside varying concentrations of sodium fluoride (NaF) showed, as expected (3), an increase in GFP fluorescence as fluoride concentrations increased up to 3 mM, followed by a decrease in fluorescence at higher concentrations (Figure 2A). This decrease is likely caused by fluoride inhibition of the gene expression machinery (40) and is consistent with previous studies (3). Accordingly, we used 3 mM NaF for the remainder of this study to obtain the expected maximum fluorescent output of the system.

**Fig. 2.**
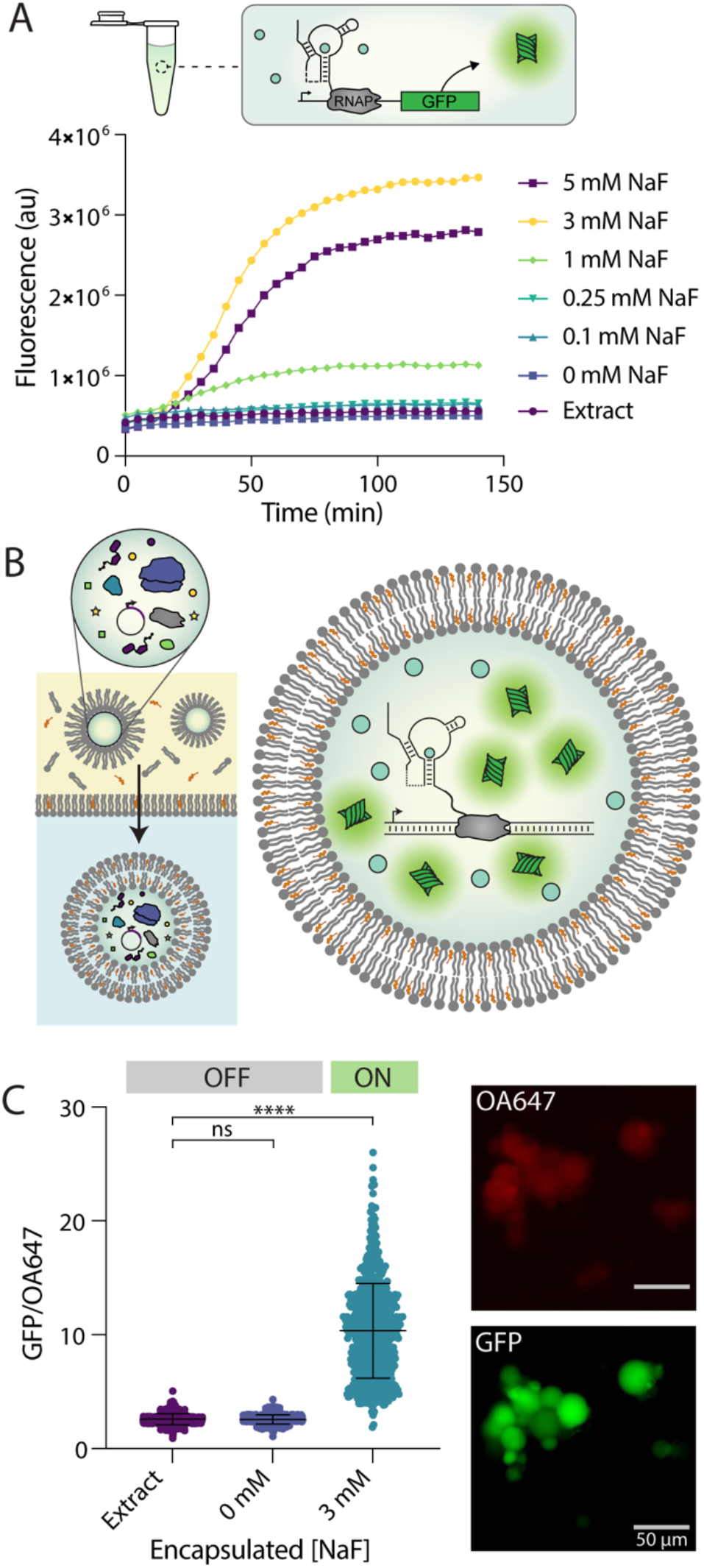
A fluoride riboswitch can function within bilayer vesicles. **A**. Riboswitch-regulated GFP expression in bulk conditions in response to increasing fluoride concentrations. In the presence of NaF the riboswitch folds into an “ON” state, which allows expression of a GFP reporter molecule. **B**. Double emulsion assembly allows the encapsulation of functional cell-free reactions. Assembled reactions are vortexed into a lipid/oil mixture, then centrifuged into an aqueous solution (left). The resulting vesicles contain cell-free reactions which can respond to co-encapsulated fluoride by expressing GFP (right). **C**. GFP/OA647 fluorescence, which indicates GFP concentration relative to the OA647 volume marker inside each liposome. GFP/OA647 fluorescence increases inside of vesicles when 3 mM NaF is co-encapsulated compared to no DNA (Extract) or no fluoride (0 mM NaF) controls. Micrographs show variations in GFP fluorescence between vesicles from the same population, which results in a distribution of fluorescence values. Scale = 50 µm. Black lines indicate mean fluorescence and standard deviation. **** p ≤ 0.0001, nonsignificant (ns) p > 0.1234; p-values generated using a One-Way ANOVA and Tukey’s Multiple Comparisons Test.

We then set out to assess whether the fluoride riboswitch could retain functionality when encapsulated within lipid vesicles. Vesicles were synthesized using a water-in-oil emulsion transfer method (Figure 2B) (41). In this method, various membrane amphiphiles (e.g. lipids, cholesterol, fatty acids, diblock copolymers) are dissolved into an oil phase and an emulsion is formed by vortexing the aqueous cell-free reaction into this mixture. The emulsion is then layered onto a second aqueous layer, and emulsified droplets are centrifuged through the oil-water interface to generate unilamellar vesicles. Vesicle synthesis using this approach yields a distribution of different vesicle sizes on the 5-50 µm scale, which could impact our quantification of fluorescence (42). To control for this, we also incorporated a protein-conjugated dye, ovalbumin-conjugated Alexafluor 647 (OA647), which served as a volume marker and allowed us to detect the vesicle interior regardless of GFP expression level (30, 43). After synthesis, vesicles were incubated under varying conditions at 37°C, and protein expression was assessed using epifluorescent microscopy. Vesicles were imaged using GFP and Cy5.5 channels, and images were analyzed using the NIS-elements AR software program (44), which allowed us to automatically select vesicle interiors using the OA647 marker and report GFP fluorescence in those regions. This protocol allowed us to analyze hundreds of vesicles per sample, maintain the same selection parameters between samples, and minimize the impact of user selection bias in the analysis. Additionally, the encapsulated volume marker allowed us to report GFP expression relative to OA647 fluorescence to control for possible variability in vesicle size or loading. Using this method, we were able to ensure that our measurements were isolated to intact (non-lysed) vesicles which retained their protein cargo (Figure S1).

Using the above approach, we encapsulated cell-free reactions with and without fluoride present in the bulk reaction mixture. We chose to use a 2:1 ratio of cholesterol and POPC phospholipid as membrane amphiphiles due to their previous use in similar encapsulated expression studies (21, 28–30, 43). Upon co-encapsulation of the riboswitch with 3 mM NaF we observed GFP expression inside vesicles, indicating the riboswitch was in the “ON” state (Figure 2C). In contrast, in the absence of DNA (extract only) or in the absence of fluoride (0 mM NaF) we observed minimal GFP expression, indicating an “OFF” state (Figure 2C). This high level of GFP induction inside vesicles by fluoride indicates that membrane encapsulation does not eliminate the ability of the riboswitch to fold properly and does not cause significant nonspecific expression.

We observed that populations of vesicles exhibited variations in GFP fluorescence between individual liposomes after 6 hours of incubation (Figure 2C), a phenomenon which has been observed in similar studies across multiple encapsulation protocols (21, 24, 30, 42, 45–47). It has been hypothesized that these variations in gene expression may be caused by variability in vesicle loading and/or varied levels of molecular exchange with the surrounding buffer for vesicles of different sizes (24, 45, 46, 48). To report this variability across vesicle populations we have included metrics of skew for each population result (Tables S1-S3). Even after taking this variability into account, however, induction of GFP expression is clearly observable across the vesicle population, indicating proper riboswitch sensor activity and a robust response to fluoride in encapsulated sensors.

### External fluoride can be detected by an encapsulated riboswitch

We next sought to determine whether the encapsulated riboswitch could detect fluoride added to the external solution of pre-assembled sensor vesicles. To assess this, we prepared vesicles containing cell-free reactions without NaF present in the reaction mixture. We then titrated in NaF into the solution surrounding vesicles (Figure 3A) and imaged vesicles following incubation for 6 hours at 37°C. We observed increasing GFP expression with increasing concentrations of NaF up to 3 mM and a slight decrease in average fluorescence at 5 mM, consistent with bulk studies (Figure 3B-D). All fluoride-containing conditions exhibited a significant increase in fluorescence compared to no-DNA and no-fluoride controls (Figure 3B & C, Table S1). When incubated with chloride, a similarly monovalent anion, a slight response to increasing ion concentration was observed, however these responses were significantly lower than any response to fluoride and did not exhibit any of the highly active vesicles that were observed in all fluoride-containing conditions (Figure S2). These responses were easily distinguishable between fluoride and chloride, indicating sufficient specificity to fluoride, as has been observed previously (3). Taken together, these results indicate that increasing concentrations of fluoride added to the extravesicular environment can be detected by the encapsulated riboswitch.

**Fig. 3.**
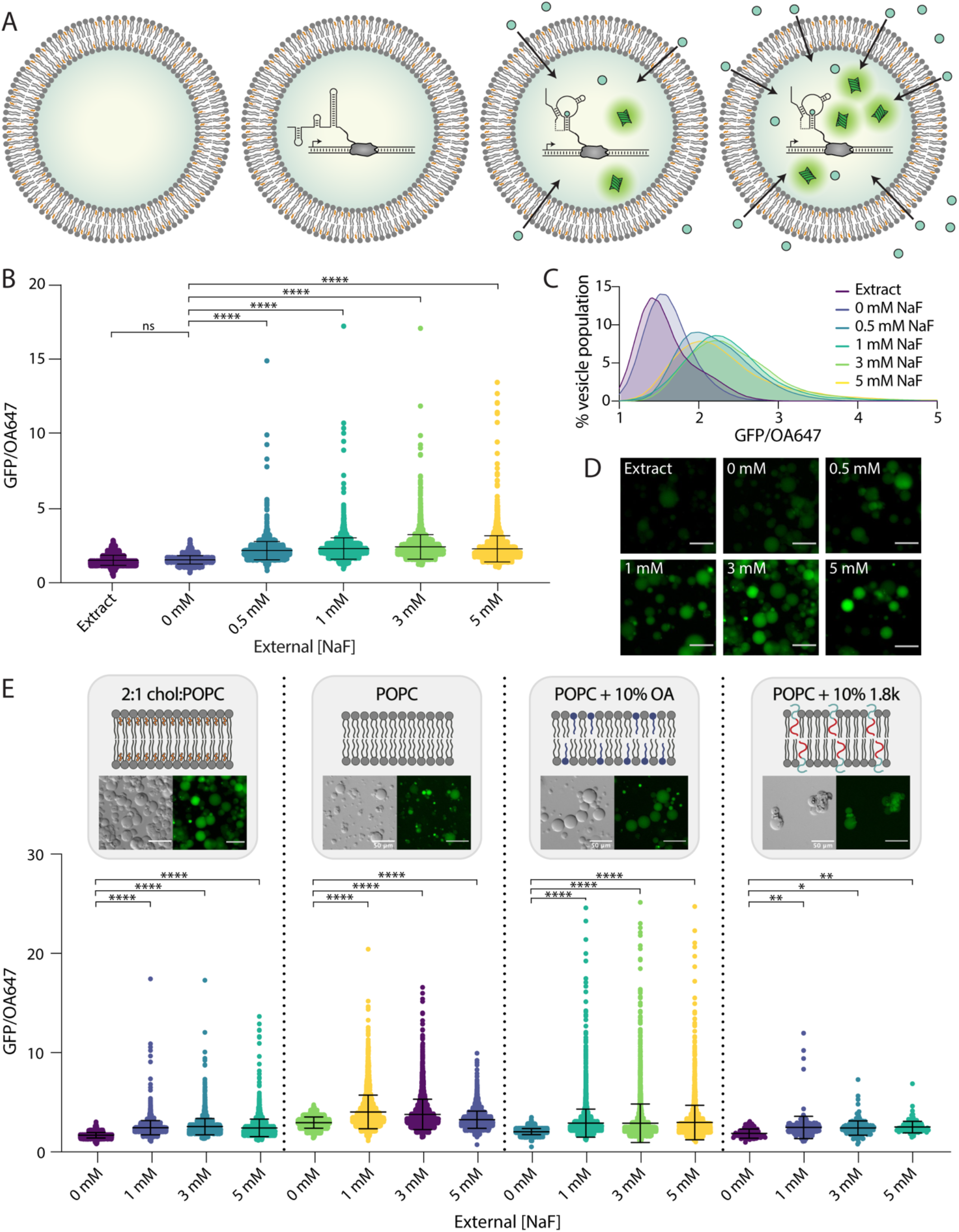
Detection of external fluoride by encapsulated sensors. **A**. Schematic of conditions. Vesicles were prepared encapsulating extract only (left), or fully assembled reactions without NaF. Upon addition of increasing fluoride in the external solution, expression of GFP inside vesicles increases (right). **B**. GFP/OA647 fluorescence as a result of riboswitch activity in 2:1 cholesterol:POPC vesicles in response to increasing NaF added externally. Black lines indicate mean fluorescence ratio and standard deviation. **C**. Histogram of vesicle populations shown in (B). Data plotted with lowless curve fitting. **D**. GFP fluorescence in micrographs of vesicles with increasing external concentrations of NaF. Scale = 50 µm. **E**. GFP/OA647 fluorescence in response to increasing fluoride shown from left to right: 2:1 cholesterol:POPC membranes (data from B); pure POPC lipid membranes; POPC + 10% oleic acid membranes; POPC + 10% 1.8k PEO-b-PBD membranes. Composition and morphology of each membrane composition indicated by schematics and micrographs, respectively. **** p ≤ 0.0001, ** p ≤ 0.0021, * p ≤ 0.0332, nonsignificant (ns) p > 0.1234; p-values generated using a One-Way ANOVA and Tukey’s Multiple Comparisons Test.

This result was somewhat unexpected, as we anticipated that the membrane would be relatively impermeable to charged fluoride ions. The observed magnitude of fluoride permeability may be explained in part by the transient formation of hydrofluoric acid (HF). HF has been shown to exhibit a permeability coefficient that is seven orders of magnitude greater than fluoride anions through lipid/cholesterol bilayers, indicating that HF travels through the membrane much more readily than its anionic F^-^ counterpart (40, 49). We confirmed this effect by encapsulating a pH sensitive dye, HPTS, which reported a slight decrease in pH in the vesicle interior upon the addition of fluoride to the external buffer (Figure S3). This result indicates an increase in proton concentration inside the vesicle as fluoride concentration increases, consistent with cross-membrane transport of HF.

Since we observed fluoride could pass through the membrane to interact with the encapsulated riboswitch, we wondered if we could alter the composition of vesicle membranes to modulate membrane permeability and thereby modulate sensitivity of these sensors to external fluoride. Membrane permeability to small molecules depends significantly on membrane composition, as various lipid chain chemistries and contributions from other amphiphilic components can impart an effect on membrane physical properties. Cholesterol, a major component of our original 2:1 cholesterol:POPC lipid composition, is known to decrease membrane permeability by increasing lipid packing and altering membrane fluidity and rigidity (50). PEO-b-PBD diblock copolymers are similarly known to reduce membrane permeability by increasing membrane viscosity, introducing steric barriers from the polyethylene glycol groups that assemble at the membrane interface, and increasing thickness and chain entanglements within the hydrophobic portions of the membrane (51, 52). In contrast, fatty acids such as oleic acid have been shown to increase membrane permeability to ionic solutes by incorporating single hydrocarbon chains that have a different shape and amphiphathicity than diacyl chains, reducing lipid chain packing and enhancing fluidity of the bilayer (53, 54). Using this series of amphiphilic molecules, we set out to assess the capacity of membrane amphiphiles and the resulting membrane permeability to modulate the performance of an encapsulated cell-free sensor.

To explore the effect of these amphiphiles on membrane permeability to fluoride, we prepared vesicles with either 1) pure POPC lipid, 2) POPC lipid + 10% oleic acid (OA), or 3) POPC lipid + 10% PEO14-b-PBD22 polymer (MW = 1.8kDa, hereafter referred to as 1.8k) components in the lipid/oil mixture, encapsulating cell-free reactions as normal (Figure 3E). We observed an increase in overall GFP expression in both pure POPC lipid and POPC + 10% OA conditions compared to our original 2:1 cholesterol:POPC lipid composition. These results are consistent with the removal of cholesterol and the addition of oleic acid, respectively, both of which should increase membrane permeability (Figure 3D). In addition, the inclusion of oleic acid in vesicle membranes led to a reduction of sensor sensitivity, measured via a reduced concentration dependence of GFP expression on NaF concentration, indicating high permeability to any amount of external fluoride. In contrast, vesicles containing 10% 1.8k diblock copolymer exhibited very little GFP expression, indicating reduced membrane permeability. Mean POPC vesicle fluorescence peaked at 1 mM NaF, while 10% OA and 10% 1.8k diblock copolymer responses were maximum at 5 mM NaF (Table S2). Taken together, these results indicate that exchanging membrane components to control membrane permeability provides a handle to tune the sensitivity of an encapsulated riboswitch to an analyte of interest. Further, the selection of highly permeable amphiphiles does not necessarily improve sensor performance and may instead increase overall signal but limit sensor resolution. A balance between analyte access and desired sensing behavior is likely an important consideration for engineering encapsulated biosensing systems depending on the desired application.

### Encapsulation protects sensor components from degradation

Having established that these vesicle sensors can detect external fluoride, we next wanted to explore how they might function in complex samples. One of the major benefits of membrane encapsulation is the ability to leverage the semipermeable barrier formed by the membrane to contain and protect encapsulated components. Cell-free reactions, particularly those using riboswitches, are highly sensitive to the presence of nucleases and proteases which can degrade sensor components before a target analyte is encountered (19). Due to their large size, however, enzymes are unable to pass through the vesicle membrane to access encapsulated reactants.

To determine whether the vesicle membrane can sufficiently protect encapsulated reactions from external degradation, we tested various vesicle assemblies in the presence of RNAse A (Figure 4A). We observed that RNAse completely eliminated the riboswitch response to NaF both in bulk conditions and when RNAse was co-encapsulated with the cell-free reaction in vesicles (Figure 4B, Figure 4C). In contrast, encapsulated sensors maintained the ability to respond to externally added NaF when RNAse was present in the external sample (Figure 4D, Table S3). Interestingly, we noticed a decrease in mean GFP fluorescence at higher external NaF concentrations compared to sensors without RNAse present, which we hypothesized was due to slightly higher degrees of vesicle instability or membrane permeability from the addition of small amounts of glycerol in the RNAse buffer (Figure S4). Instability could lead to higher rates of vesicle lysis and therefore lower overall GFP fluorescence, while increased permeability could cause increased reaction poisoning with high fluoride concentrations. Nevertheless, all vesicle populations exhibited increased GFP expression in the presence of fluoride, demonstrating simultaneous permeation of fluoride into the vesicle interior and exclusion of RNAse A from the cell-free reaction.

**Fig. 4.**
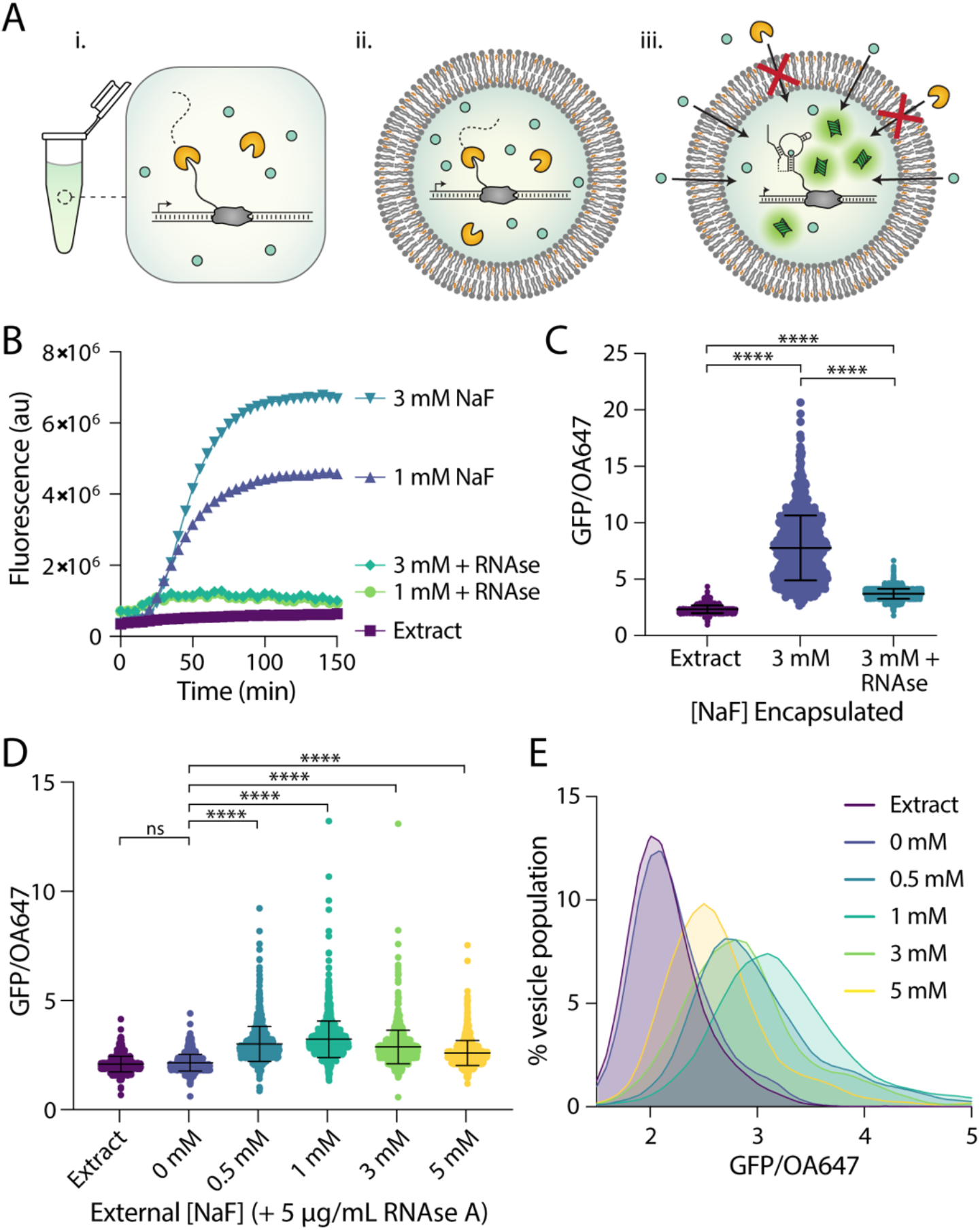
Encapsulation protects from degradation by RNAse A. **A**. Schematic of RNAse-containing conditions. RNAse A degrades the riboswitch **(i)** in bulk conditions and **(ii)** when co-encapsulated with reactants but is unable to reach reactants contained within vesicles **(iii). B**. Riboswitch response to NaF in bulk conditions with and without RNAse A added to reaction. **C**. Riboswitch activity as indicated by GFP/OA647 fluorescence when encapsulated with 3 mM NaF compared to the co-encapsulation of both 3 mM NaF and RNAse A. **D**. Response of encapsulated riboswitch to externally added NaF with RNAse A present in external solution. Black lines indicate mean and standard deviation. **E**. Histogram of data in (D). Data plotted with lowless curve fitting. **** p ≤ 0.0001, nonsignificant (ns) p > 0.1234; p-values generated using a One-Way ANOVA and Tukey’s Multiple Comparisons Test.

### Vesicle-based sensors can incorporate a colorimetric readout and detect fluoride in real-world samples

Finally, we wondered if we could extend these results to conditions that would be more relevant for real-world environmental sensing. Although fluorescence is a common readout for many biological assays, GPF fluorescence inside vesicles is difficult to monitor with common equipment, particularly in non-laboratory settings. To address this limitation, we coupled fluoride detection to an alternative reporter enzyme, catechol (2,3)-dioxygenase (C23DO) (3). In this system, the riboswitch “ON” state leads to the expression of C23DO, which catalyzes the conversion of its colorless substrate, catechol, to the yellow-colored 2-hydroxymuconate semialdehyde to generate a colorimetric response (Figure 5A). In bulk conditions this construct exhibits a fast and robust response to fluoride, and the colorimetric output generated is clearly distinguishable by eye for both laboratory and field-collected water samples (3).

**Fig. 5.**
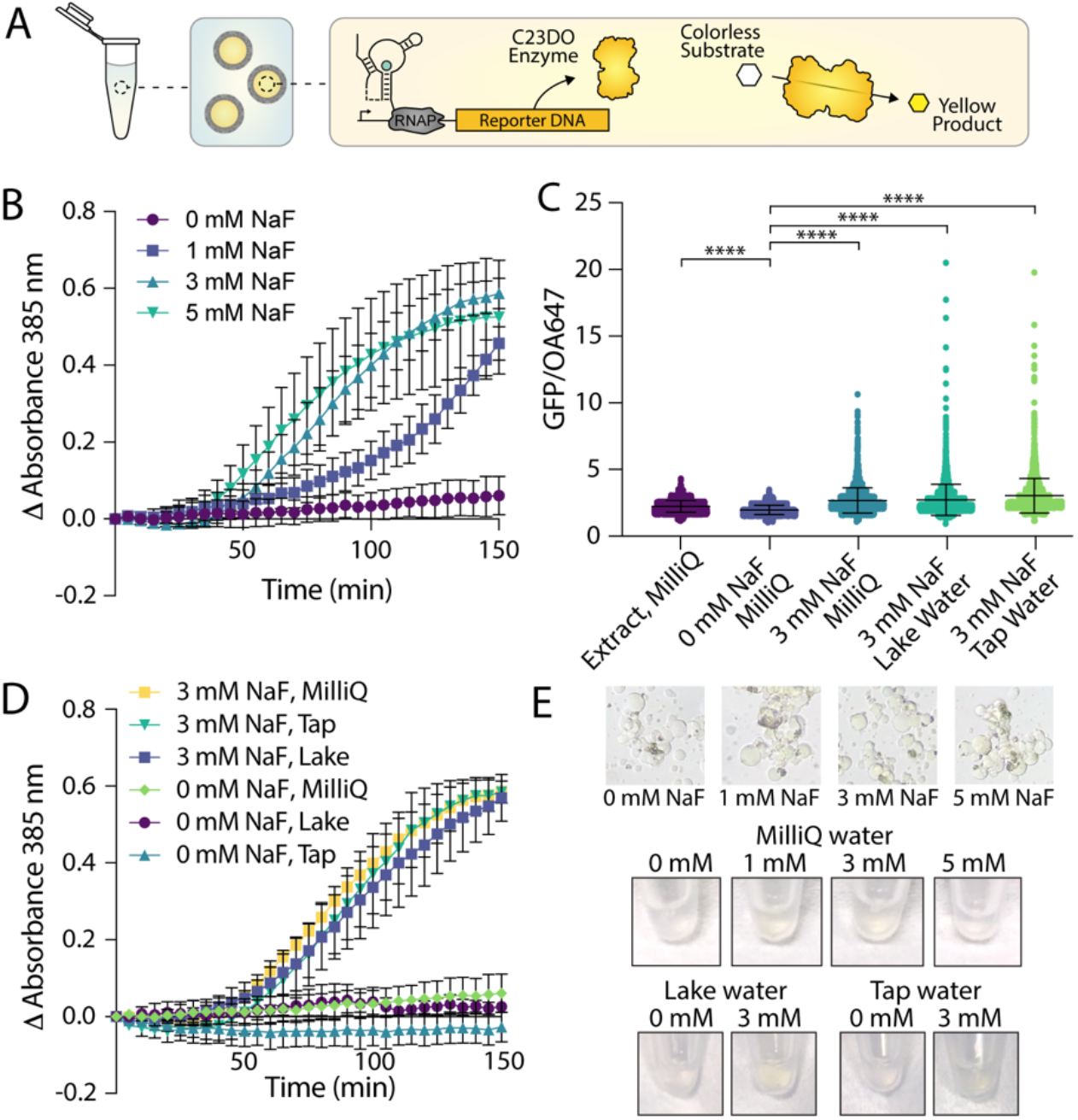
Enzymatic readout and detection of fluoride in real-world samples. **A**. Schematic of encapsulated enzymatic readout. Riboswitch activation inside vesicles leads to the expression of C23DO, resulting in the production of a yellow product that is localized to the vesicle interior. **B**. Absorbance over time inside of vesicles encapsulating a catecholase-based readout in response to external NaF. Output is reported as a change in absorbance to account for variations in final vesicle concentration between different vesicle preparations. N=3 independent vesicle preparations. **C**. GFP/OA647 fluorescence ratios observed in vesicles containing a GFP-based readout after incubation in outer solutions from laboratory grade water (MilliQ), tap water, and lake water supplemented with NaF. **D**. Absorbance over time inside of vesicles incubated in samples derived from MilliQ water, tap water and lake water supplemented with either 0 mM NaF or 3 mM NaF. N=2 independent vesicle preparations. **E**. Colorimetric changes in vesicles as viewed through a microscope eyepiece and by eye in Eppendorf tubes. **** p ≤ 0.0001, nonsignificant (ns) p > 0.1234; p-values generated using a One-Way ANOVA and Tukey’s Multiple Comparisons Test.

To investigate whether this enzymatic reporter could function within our sensor vesicles, we encapsulated cell-free reactions with DNA coding for the riboswitch-C23DO construct and supplemented them with 1 mM catechol (Figure 5A). We then titrated NaF into the outer solution and monitored color changes in each population of vesicles via changes in absorbance at 385 nm. In contrast to our GFP-based readout, signal amplification from the enzyme-regulated construct allowed us to assess absorbance changes in an entire population of vesicles rather than on a vesicle-by-vesicle basis. RNAse A was also added to the outer vesicle solution to control for any unencapsulated reactions caused by vesicle lysis. The response was significantly lower than that observed in bulk (Figure S5), but increases in absorption in response to increasing NaF concentrations were observed across multiple sample preparations (Figure 5B). Readout time plays a key role in sensor response (3), particularly for the 1 mM NaF condition, where amplified responses followed by signal decay make quantification difficult (Figure 5B, Figure S5). Little expression was observed in 0 mM NaF samples in this time frame, however, which indicates potential for these sensors to serve as binary classifiers even in the presence of low fluoride concentrations. Importantly, responses to fluoride could be detected within 2 hours of incubation compared to 6 hours for the GFP-based readout.

Having demonstrated the compatibility of our encapsulated sensors with multiple reporters and their ability to detect extravesicular fluoride, we set out to test whether these sensors could be used to monitor fluoride concentrations in real-world samples. We collected water samples from Lake Michigan and the Evanston, IL municipal tap water supply and used each sample to prepare the vesicle outer solution. We supplemented these outer solutions with either 0 mM NaF or 3 mM NaF and added vesicles with either the GFP-coupled riboswitch construct or the colorimetric construct. We observed increased GFP expression in all populations of vesicles incubated with 3 mM NaF outside compared to no-fluoride controls, with a slightly higher level of GFP expression in both lake and tap water samples compared to those incubated with laboratory-grade Milli Q water (Figure 5C). Similarly, vesicles encapsulating the enzymatic construct showed increasing absorption over time in the presence of 3 mM NaF, while all no-fluoride controls exhibited no significant changes in absorption (Figure 5D). Slight color changes were visible by eye in tubes containing vesicles, and changes in color inside of vesicles could be observed on the microscope as imaged through the eyepiece (Figure 5E). These results were consistent with those observed in bulk assays, which showed a slightly higher response to both tap and lake water and a significant difference between all 0 mM and 3 mM conditions (Figure S5). The results observed here highlight the feasibility of these vesicle-based sensors to detect environmentally relevant small molecules in real-world samples, a step toward encapsulation to generate deployable cell-free sensors.

## Discussion

To our knowledge, this work represents the first demonstrated function of a transcriptionally regulated riboswitch encapsulated in bilayer vesicles. We have demonstrated that this encapsulated riboswitch can detect exogenous fluoride through permeability-based sensing, generating both fluorescent and colorimetric outputs. Additionally, we have shown that responses to fluoride can be modulated by changing membrane composition, which provides a useful handle to control sensor stability and sensitivity. Looking ahead toward sensor deployment, this work establishes that encapsulation can protect cell-free sensors from degradative sample components while allowing analyte detection in real-world samples. While cell-free sensors have been previously used for the detection of environmental molecules of interest (1–9), encapsulation of these systems may ultimately diversify the contexts within which cell-free sensors can operate.

Although encapsulation can provide powerful advantages to cell-free sensing, it also brings some limitations. The concentrations of fluoride assessed here are high compared to the Maximum Contaminant Limits set by the Environmental Protection Agency (0.5 mM vs 0.22 mM (55)), which were chosen based on the spread of responses observed in liposomes. The variability observed in the responses of individual liposomes within a vesicle population would likely serve as a hurdle for technological use of these sensors in future applications, which may necessitate alternative vesicle assembly techniques, such as microfluidics (42), and a better understanding of the underlying biophysics of cell-free reactions inside membranes. While we explored protein-based outputs here, riboswitch expression could also be coupled to transcription-based reporting, such as aptamer-dye outputs (3), to build a transcription-only sensor that would require encapsulation of fewer components and should operate on quicker timescales. Finally, the reintroduction of a barrier between sample and sensor also requires strategies to transport specific analytes into the vesicle. Membrane compositional changes can enable permeability-based import for certain small analytes, with many natural and synthetic amphiphiles to choose from. Moving forward, we can gain even finer control of membrane permeability by incorporating transmembrane proteins to enhance sensing capabilities and introduce more advanced sensing or responsive functions. These strategies could ultimately allow new functions for these types of sensors, including conjugation-based capture methods, deployment and transport of cell-free reactions, controlled sensor degradation, or enhanced sensor biocompatibility (18).

The diversity of existing cell-free sensors could ultimately lead to a new generation of encapsulated biosensors for a wide array of analytes. With the modularity of components in these systems, vesicle-based sensors could be engineered which use various membrane components, genetic circuits, and triggered responses to detect small molecules of interest (30, 56, 57). As focus shifts toward sensor application, these platforms could offer additional handles with which to tune sensor characteristics to advance the types of contexts in which cell-free sensing can operate, allowing for detection in environments like soil, ground water, or biological samples. Finally, the incorporation of additional transcription-based cell-free systems, particularly those using riboswitch-based sensing, may ultimately allow the development of a family of encapsulated sensors that are fast, specific, and deployable.

## Materials and Methods

### Chemicals

POPC (1-palmitoyl-2-oleoyl-glycero-3-phosphocholine) and cholesterol were purchased from Avanti Polar Lipids Inc. Oleic acid (OA), glycerol, sucrose, glucose, HPTS (8-Hydroxypyrene-1,3,6-trisulfonic acid trisodium salt), BioUltra Mineral Oil, phosphate-buffered saline (PBS), bovine serum albumin (BSA), and NaF were purchased from Millipore Sigma. 1.8k PEO-b-PBD polymer was purchased from Polymer Source. Ovalbumin-conjugated AlexaFluor 647 (OA647), calcein dye and HEPES buffer were purchased from Thermo Fisher. RNAse A was purchased from New England Biolabs.

### Plasmid construction

Plasmids were assembled using Gibson assembly (New England Biolabs, Cat#E2611S) and purified using a Qiagen QIAfilter Midiprep Kit (QIAGEN, Cat#12143). Coding sequences of the plasmids consist of the crcB fluoride riboswitch from Bacillus cereus regulating either superfolder GFP (pJBL3752) or catechol 2,3-dioxygenase (pJBL7025), all expressed under constitutive Anderson promoter J23119. Plasmid sequences available on Addgene with accession numbers 128809 (pJBL3752) and 128810 (pJBL7025).

### Cell-free reaction assembly

Cell-free extract and reactions were prepared according to established protocols (3, 39). Briefly, cell-free reactions were assembled by mixing cell extract, a reaction buffer containing the small molecules required for transcription and translation (NTPs, amino acids, buffering salts, crowding agents, and an energy source), and DNA templates and inducers at a ratio of approximately 30/30/40. Sucrose was added to a final concentration of 200 mM to facilitate encapsulation. Each reaction was prepared on ice to 16.5 µL final volume in batches of 7. Reactions were prepared with 10 nM pJBL3752 (riboswitch-GFP plasmid) or pJBL7025 (riboswitch-enzyme plasmid) + 1 mM catechol. Reaction master mix was assembled, then added to DNA, inducers, sucrose, and water to a final volume of 16.5 µL per reaction aliquot. For reactions containing volume marker, reaction mix was supplemented with 1.4 µL OA647. Preparation conditions were kept consistent between reactions, only varying NaF concentration or omitting DNA for extract-only controls.

### Encapsulation of cell-free reactions

Encapsulated sensors were prepared via water-in-oil double emulsion methods. Lipid films were prepared by mixing amphiphiles (lipid, cholesterol, fatty acid or polymers) in chloroform to a final amphiphile concentration of 25 mM at a volume of 200 µL. Films were dried onto the side of a glass vial under nitrogen gas, then placed in a vacuum oven overnight. 200 µL of BioUltra mineral oil was added to lipid films and heated at 80°C for 30 minutes, followed by 10 seconds of vortexing to incorporate amphiphiles into the oil. Lipid/oil mixtures were cooled on to room temperature, then placed on ice during cell-free reaction assembly. Cell-free reactions were prepared on ice as described above. Reactions were layered on top of lipid/oil mixture, then vortexed for 30 seconds to form an emulsion. Emulsions were incubated at 4°C for 5 minutes, then layered onto outer solution containing all small molecules required for transcription and translation, 100 mM HEPES buffer (pH 8), and 200 mM glucose. Samples were again incubated at 4°C for 5 minutes, then centrifuged for 15 minutes at 18,000 rcf at 4°C. Vesicle pellets were collected by pipette and placed into fresh Eppendorf tubes. Prepared vesicles were then added in 10 µL aliquots to 20 µL fresh outer solution supplemented with NaF, certain water samples and/or RNAse A (5 µg/mL final concentration). Osmolarity of NaF stock solution was adjusted to match that of the outer solution by adding glucose to minimize osmotic effects on vesicles.

### Cell-free protein expression

For bulk assays, unencapsulated reactions were prepared as described above and added to 384-well plates. Protein expression was monitored at 37°C in a SpectraMax i3x plate reader (Molecular Devices). GFP was monitored at ex: 485 nm, em: 510 nm. Catechol absorbance was monitored at 385 nm.

Encapsulated sensors with a colorimetric readout were monitored at an absorbance of 385 nm using a SpectraMax i3x plate reader at 37°C until expression reached a plateau, about 2.5 hours, after which samples were removed from plates and placed into Eppendorf tubes or microscopy chambers for imaging. Images of tubes and through the microscope eyepiece were taken using an iPhone 8. Absorbance measurements in the plate reader are reported relative to initial absorbance to control for slight differences in vesicle concentration between vesicle preparations.

Encapsulated sensors expressing GFP were incubated in outer solution for 6 hours at 37°C, then imaged on a Nikon Ti2 inverted microscope. Imaging chambers were blocked with BSA for 20 minutes, then triple rinsed with 766 mOsm PBS. Vesicles were added to equiosmolar PBS and allowed to settle for 5 minutes before imaging. Images were taken using DIC, GFP (ex: 470, em: 525) and Cy 5.5 (ex: 650, em: 720) filters under 10x magnification, 20% laser intensity, and 1 second exposure. Images were analyzed using Nikon NIS-elements AR software Advanced Analysis tool (44): vesicles were selected using the OA647 channel. General analysis protocol was set with the following settings. Preprocessing: Local contrast, size 105, power 50%. Threshold minimum: 393. Smooth 1x, clean 1x. Size minimum: 2 µm. Return Mean GFP, Mean OA647, Max GFP.

### Encapsulated dye assays

HPTS assay: vesicles were prepared via thin film hydration with 33% Cholesterol and 66% POPC. Lipid and cholesterol in chloroform were dried onto the side of a glass vial under nitrogen gas to form a lipid film. Vesicle films were hydrated with HEPES + 0.5 mM HPTS dye overnight at 60°C. Vesicles were extruded to 400 nm, purified via Size Exclusion Chromatography (SEC), and added to a 384-well plate with equiosmolar HEPES buffer + varying concentrations of NaCl and NaF. HPTS fluorescence was monitored with excitation at 405 and 450 nm and emission at 510 nm, as characterized by Hilburger et al. (59). HPTS fluorescence is reported as the ratio of emission intensities when excited at 450 nm/405 nm.

Calcein assay: vesicles were prepared via thin film hydration with 33% Cholesterol and 66% POPC. Lipid and cholesterol in chloroform were dried onto the side of a glass vial under nitrogen gas to form a lipid film. Vesicle films were hydrated with HEPES + 20 mM calcein dye overnight at 60°C. Vesicles were extruded to 400 nm, purified via SEC, and added to a 384-well plate with equiosmolar HEPES buffer and increasing volumes of 0.02% glycerol solution or RNAse prepared in buffer to the same final glycerol concentration (1.25 µL corresponds to the concentration used for manuscript studies). Vesicles were incubated for 4 hours at 37°C and calcein fluorescence was measured (ex: 495 nm, em: 515 nm). Vesicles were lysed with 1 µL 10% TritonX and total calcein fluorescence was measured to determine fraction release.

### Statistical Analysis

All graphing and statistical analysis was conducted in Graphpad Prism (58). Populations of vesicles were analyzed using One-Way ANOVA analysis with Tukey’s Multiple Comparisons Test and descriptive statistics. Significance is reported as follows: **** p ≤ 0.0001, ** p ≤ 0.0021, * p ≤ 0.0332, nonsignificant (ns) p > 0.1234.

## Acknowledgments

We thank Zoila Jurado and the Murray Lab as well as the Build a Cell community for providing a water-in-oil vesicle formation protocol which we adapted for these studies. We would also like to thank Adam Silverman and Dylan Brown for preparing cell extract and reaction materials.

## Funding

Northwestern University Chemistry of Life Processes Institute Cornew Innovation Award (NPK, JBL)

National Science Foundation grant 1844219 (NPK)

National Science Foundation grant 1844336 (NPK, JBL)

National Science Foundation grant 2145050 (NPK)

U.S. Department of Defense National Science & Engineering Graduate Fellowship (MAB)

## Author contributions

Conceptualization: MAB, WT, JBL, NPK

Methodology: MAB, WT, JBL, NPK

Investigation: MAB, WT

Data Analysis: MAB, WT, JBL, NPK

Visualization: MAB

Writing—original draft: MAB, NPK

Writing—review & editing: MAB, WT, NPK, JBL

## Competing interests

Authors declare that they have no competing interests.

## Data and materials availability

Plasmid sequences available on Addgene with accession numbers 128809 (pJBL3752) and 128810 (pJBL7025).

## Notes

### Competing Interest Statement

The authors have declared no competing interest.

