## Supporting Information for "Robust and tunable performance of a cell-free biosensor encapsulated in lipid vesicles"

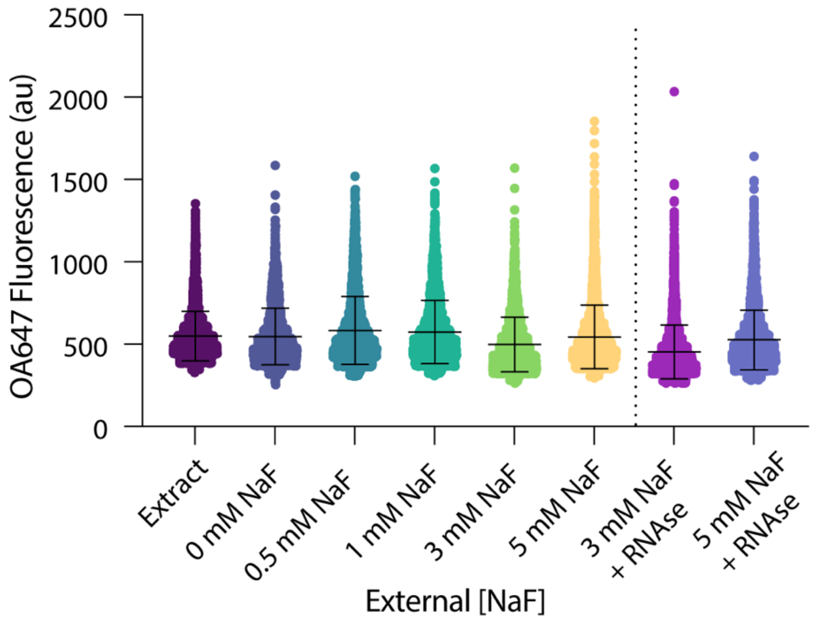


**Fig. S1. OA647 retention in 2:1 cholesterol:POPC vesicles following encapsulation and protein expression.**

Vesicle populations exposed to increasing fluoride in the external solution exhibit the retention of a volume marker, OA647, even when RNAse A is present externally. While average fluorescence varies slightly between populations, corresponding to differences in the size of analyzed vesicles between conditions, all samples exhibit similar fluorescence profiles consistent with the retention of protein-sized molecules within the vesicle interior.

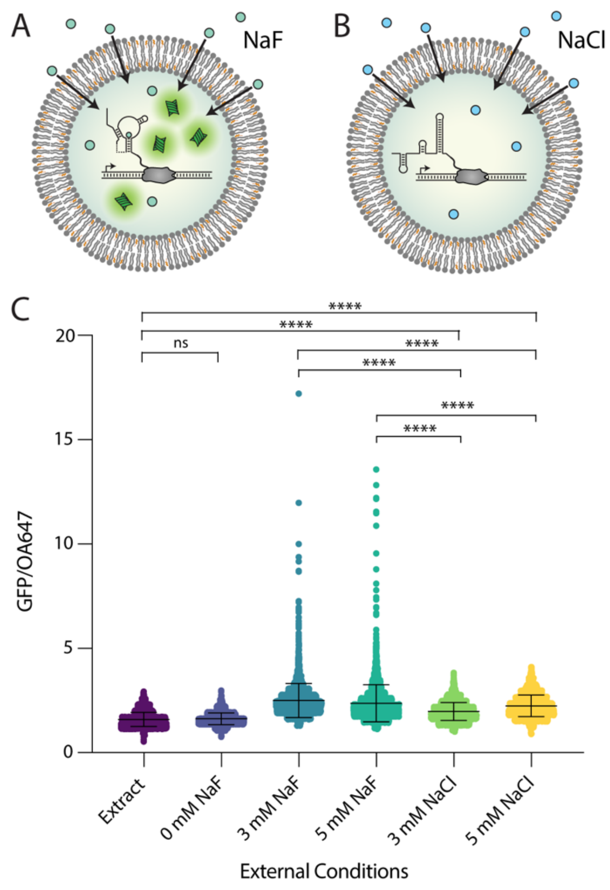


**Fig. S2. An encapsulated riboswitch responds specifically to fluoride.**

Fluoride permeates the vesicle membrane to initiate the expression of a GFP reporter inside vesicles. B. The external addition of NaCl does not result in robust GFP expression inside vesicles. C. GPF/OA647 fluorescence in vesicles with either NaF or NaCl added to the external buffer. While a response is observed to increasing chloride, the magnitude is significantly less than the response to fluoride and there is no observed population shift towards highly active vesicles. Differences in expression between fluoride and chloride containing conditions were clearly distinguishable, indicating sufficient specificity to fluoride over chloride. **** $p \leq 0.0001$, nonsignificant (ns) $p > 0.1234$; p-values generated using a One-Way ANOVA and Tukey's Multiple Comparisons Test.

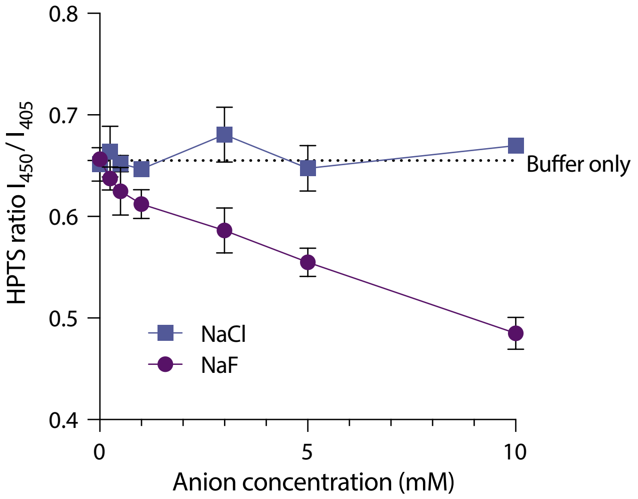


**Fig. S3. pH decreases as increasing NaF is added externally to vesicles.**

Lipid/cholesterol vesicles containing HPTS dye without cell-free expression systems show changes in fluorescence after addition of anions to the external solution, indicating a cross-membrane effect on pH caused by externally added NaF. Compared to NaCl and buffer only controls, the pH of the vesicle interior decreases in the presence of externally added NaF, as indicated by a decreasing fluorescence ratio of HPTS, a pH-sensitive dye. These results indicate that fluoride ions may permeate the membrane as HF, bringing $H^+$ ions with them as they pass through the membrane.

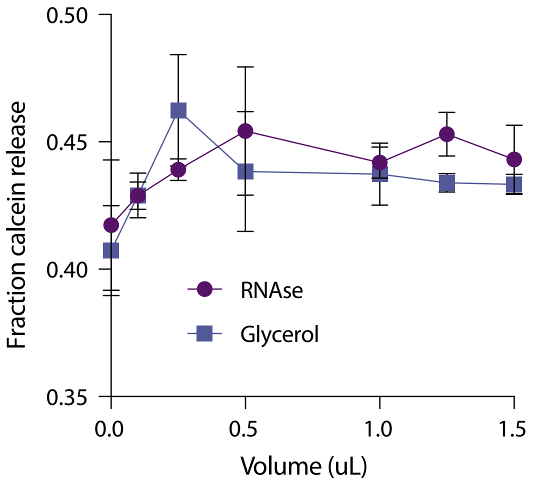


**Fig. S4. Glycerol addition increases membrane permeability.**

Lipid/cholesterol vesicles encapsulating calcein, a self-quenching fluorescent dye, show slightly increasing cargo leakage following glycerol addition to the surrounding buffer. In both the presence and absence of RNAse, addition of increasing volumes of 0.02% glycerol solutions (1.25 μL of which was added in vesicle studies) leads to slightly higher levels of calcein dye release from the vesicle interior, indicating increased membrane permeability to small molecules.

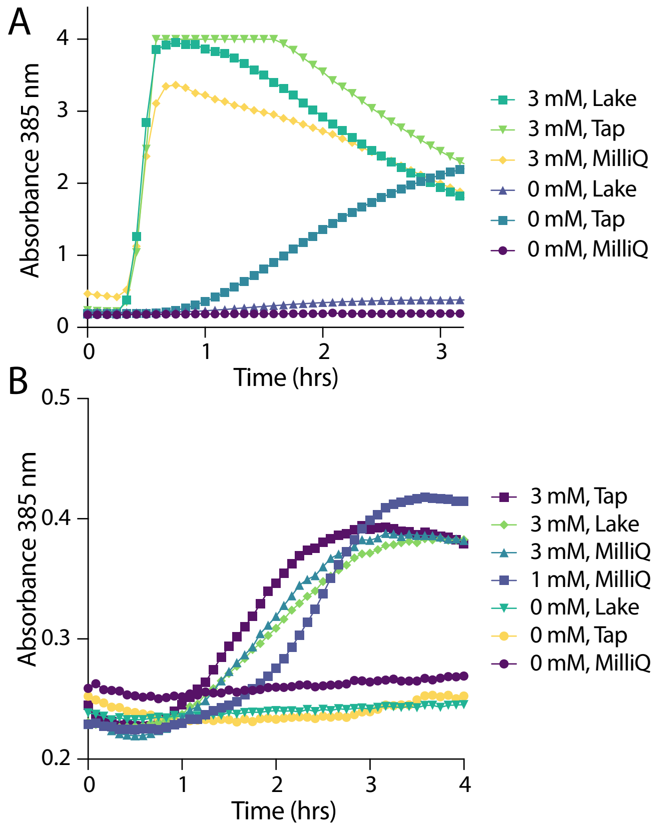


**Fig. S5. Catecholase conversion in response to fluoride in bulk and inside of vesicles.**

**A.** Bulk reactions show slightly higher responses to 3 mM NaF supplemented in water samples taken from Lake and Tap water compared to laboratory-grade MilliQ water. Absorbance was also observed to increase in unsupplemented tap water, likely due to low levels of fluoride added to public drinking supply. **B.** In vesicles, absorbance increases specifically in the presence of supplemented NaF. Quantification is difficult, with 1 mM NaF exhibiting a slightly delayed response compared to 3 mM NaF but a similar expression profile and maximum absorbance. Responses are similar between all 3 mM samples regardless of water source, with a slight increase in expression in unsupplemented tap water at later timepoints (consistent with bulk data). (n=1 example vesicle preparation shown here)

| External NaF concentration | Mean Fluorescence GFP/OA647 | SEM | Significantly different than 0 mM? | Skewness | Kurtosis |
|---|---|---|---|---|---|
| **Extract** | 1.596 | 0.0054 | ns | 0.5225 | 0.8674 |
| **0 mM** | 1.625 | 0.0062 | | 0.4007 | 0.4155 |
| **0.5 mM** | 2.250 | 0.0126 | **** | 2.205 | 13.12 |
| **1 mM** | 2.388 | 0.0143 | **** | 1.824 | 9.110 |
| **3 mM** | 2.500 | 0.0167 | **** | 2.083 | 8.960 |
| **5 mM** | 2.372 | 0.0166 | **** | 4.587 | 46.19 |

**** $p \leq 0.0001$, nonsignificant (ns) $p > 0.1234$; p-values generated using a One-Way ANOVA and Tukey's Multiple Comparisons Test.

**Table S1**: **GFP expression in response to externally added NaF in 2:1 cholesterol:POPC vesicles.**

Descriptive statistics of GFP expression in populations of vesicles exposed to increasing amounts of externally added NaF. Statistical analysis was computed compared to 0 mM NaF conditions.

| External NaF concentration | Mean GFP/OA647 | SEM | Different than 0 mM? | Skewness | Kurtosis |
|---|---|---|---|---|---|
| **POPC, 0 mM** | 2.904 | 0.01074 | | 0.1471 | -0.3680 |
| **POPC, 1 mM** | 3.968 | 0.02997 | **** | 2.063 | 7.183 |
| **POPC, 3 mM** | 3.724 | 0.02570 | **** | 2.433 | 9.854 |
| **POPC, 5 mM** | 3.190 | 0.01478 | **** | 1.528 | 5.072 |
| **10% OA, 0 mM** | 1.997 | 0.004865 | | 0.3135 | 0.1434 |
| **10% OA, 1 mM** | 2.850 | 0.02232 | **** | 6.120 | 59.59 |
| **10% OA, 3 mM** | 2.839 | 0.03147 | **** | 4.848 | 32.92 |
| **10% OA, 5 mM** | 2.909 | 0.03352 | **** | 5.038 | 38.90 |
| **10% 1.8k, 0 mM** | 1.807 | 0.05333 | | 0.6967 | -0.2151 |
| **10% 1.8k, 1 mM** | 2.414 | 0.07304 | ** | 5.698 | 39.98 |
| **10% 1.8k, 3 mM** | 2.353 | 0.04787 | * | 2.199 | 11.16 |
| **10% 1.8k, 5 mM** | 2.458 | 0.04654 | ** | 3.247 | 19.66 |

**** $p \leq 0.0001$, ** $p \leq 0.0021$, * $p \leq 0.0332$, nonsignificant (ns) $p > 0.1234$; p-values generated using a One-Way ANOVA and Tukey's Multiple Comparisons Test.

**Table S2**: **GFP expression in response to externally added NaF in vesicles with varying membrane compositions.**

Descriptive statistics of GFP expression in populations of vesicles with varying membrane compositions in response to increasing concentrations of externally added NaF. Statistical analysis was computed compared to 0 mM NaF conditions for each respective membrane composition.

| External NaF concentration | Mean Fluorescence GFP/OA647 | SEM | Significantly different than 0 mM? | Skewness | Kurtosis |
|---|---|---|---|---|---|
| **Extract only** | 2.152 | 0.013 | ns | 0.9806 | 2.661 |
| **0 mM** | 2.213 | 0.013 | | 0.9952 | 2.501 |
| **0.5 mM** | 3.071 | 0.026 | **** | 2.236 | 9.502 |
| **1 mM** | 3.289 | 0.022 | **** | 3.289 | 24.83 |
| **3 mM** | 2.930 | 0.023 | **** | 3.746 | 34.26 |
| **5 mM** | 2.662 | 0.018 | **** | 2.251 | 10.79 |

**** $p \leq 0.0001$, nonsignificant (ns) $p > 0.1234$; p-values generated using a One-Way ANOVA and Tukey's Multiple Comparisons Test.

**Table S3**: **GFP expression in response to externally added NaF in 2:1 cholesterol:POPC vesicles with RNAse A in the surrounding solution.**

Descriptive statistics of GFP expression in populations of vesicles with RNAse present in the surrounding solution in response to increasing concentrations of externally added NaF. Statistical analysis was computed compared to 0 mM NaF conditions.
